# Heartbeat-evoked potentials during interoceptive-exteroceptive integration are not consistent with precision-weighting

**DOI:** 10.1101/2021.02.03.429610

**Authors:** Leah Banellis, Damian Cruse

## Abstract

Interoceptive-exteroceptive integration is fundamental for a unified interactive experience of the world with the body. Predictive coding accounts propose that these integrated signals operate predictively, with regulation by precision-weighting. Heartbeat-evoked potentials (HEPs) are one means to investigate integrated processing. In a previous study, consistent with predictive coding characterisations of precision-weighting, we observed modulation of HEPs by attention. However, we found no evidence of HEP modulation by participants’ interoceptive ability, despite the characterisation by predictive coding theories of trait abilities as a similar reflection of differential precision-weighting. In this study, we sought to more sensitively test the hypothesised trait-precision influences on HEPs by using an individually-adjusted measure of interoceptive performance. However, contrary to a precision-weighted predictive coding framework, we failed to find evidence in support of the HEP modulations by attentional-precision or trait-precision. Nonetheless, we observed robust HEP effects indicative of an expectation of a sound on the basis of a heartbeat -i.e. interoceptive-exteroceptive integration. It is possible that under our more individually-tailored task, participants relied less on attentional-precision to ‘boost’ predictions due to an enhanced perception of cardio-audio synchrony. Furthermore, assessing interoceptive ability is challenging, thus variations in performance may not accurately reflect trait-precision variations. Nevertheless, in sum, our findings are inconsistent with a precision-weighted prediction error view of the HEP, and highlight the need for clearer definitions of the manipulation and measurement of precision in predictive coding. Finally, our robust interoceptive-exteroceptive integration HEP effects may provide a valuable tool for investigating such integration in both clinical conditions and cognition.

**Impact statement:** We investigate heart-evoked potentials during interoceptive-exteroceptive integration to determine whether cross-modal integrated processes operate under a precision-weighted predictive coding framework. Using a more sensitive individually-tailored task, we found no evidence of the modulation of cardio-audio expectation by attention or individual differences in interoceptive perception (i.e. by state or trait measures of precision). Nonetheless, we replicate evidence of cardiac-driven predictions of auditory stimuli, providing a potential tool for investigating their relationship with emotion and embodied selfhood.

---

Predictive coding accounts describe the brain as probabilistically inferring the causes of upcoming sensory events (Friston, 2010; Rao & Ballard, 1999). Under these accounts, predictions from generative models are compared with inputted sensory information, with the discrepancy computed as prediction error. Predictive mechanisms are accomplished hierarchically, with predictions feeding into each layer top-down, and prediction errors bottom-up. The predictive coding framework is linked to many aspects of cognition, perception, and action, with ‘successful’ processes resulting from the minimisation of prediction error across all levels of the hierarchy (Enns & Lleras, 2008; Wolpert & Flanagan, 2001; Kilner et al., 2007; DeLong et al., 2005; Frith & Frith, 2006; Clark, 2013; Friston, 2010).

Although initially applied to exteroceptive processing, it became apparent that for the framework to encompass the integrated experience of perceiving and interacting with the world via the body, inferences of both internal and external systems must be intertwined (Allen & Friston, 2018; Friston, 2009; Petzschner et al., 2017; Pezzulo, 2014; Seth & Friston, 2016). Thus, predictive coding models emerged that encompassed the inferential processing of the body - specifically, interoceptive signals reflecting visceral bodily sensations and internal bodily states (Barrett & Simmons, 2015; Cameron, 2002; Seth, 2013; Seth et al., 2012; Seth & Friston, 2016; Sherrington, 1952).

One method of measuring cortical interoceptive processing is via heartbeat-evoked potentials (HEPs) - averaged neural electrophysiological signals time-locked to heartbeats (Park & Blanke, 2019; Pollatos & Schandry, 2004). Although discovered more than 30 years ago (Schandry et al., 1986), the study of HEPs is still in its infancy, with debate over the appropriate pre-processing/analysis methods, and controls for correcting confounds such as the cardiac field artefact (CFA) (Coll et al., 2021; Park & Blanke, 2019). This may in part explain the diverse spatial and temporal observation of the HEP across the literature. However, it is likely that the HEP reflects contributions from multiple sources, including baroreceptors, cardiac afferents, cutaneous receptor somatosensory mapping, and neurovascular coupling (Park & Blanke, 2019).

HEPs are primarily recorded from superficial pyramidal neurons via M/EEG (i.e. the proposed location of prediction error units), therefore some have interpreted HEP amplitude to reflect precision-weighted prediction error of each heartbeat (Ainley et al., 2016; Petzschner et al., 2019). Precision is the weight given to predictions and subsequent errors, reflected by the inverse of the variance, or the uncertainty. Attention is thought to increase the precision of the prediction errors of the attended sensory channel via synaptic gain control, enhancing model updating (Friston, 2009; Hohwy, 2012). Consistent with the role of attentional precision in modulating predictive mechanisms, previous research demonstrated attentional modulation of HEPs, supporting its interpretation as a precision-weighted prediction error signal (Banellis & Cruse, 2020; Mai et al., 2018; Montoya et al., 1993; Petzschner et al., 2019; Villena-González et al., 2017; Yuan et al., 2007).

Trait variations in uncertainty, such as individual differences in the ability to accurately sense the heartbeat, are proposed to similarly modulate predictive mechanisms via precision-weighting. For example, those with high heartbeat detection performance demonstrate larger HEP amplitudes than low heartbeat perceivers, comparable to internal/external attention contrasts (Katkin et al., 1991; Pollatos et al., 2005; Pollatos & Schandry, 2004; Schandry et al., 1986). However, on the surface, this result appears at odds with a prediction error interpretation as one might expect largest HEPs in circumstances of highest error, i.e., for low heartbeat perceivers. This disparity is often reconciled via appeal to precision-weighting, such that a small prediction error weighted by high precision may result in a larger evoked potential than a large prediction error weighted by low precision (Kok et al., 2012). However, care should be taken when interpreting trait variations as different heartbeat detection tasks assess distinct processes and some tasks may not validly measure ability (Brener & Ring, 2016; Corneille et al., 2020; Desmedt et al., 2020; Ring & Brener, 2018).

The cross-modal predictive mechanisms proposed to underlie an integrated experience of perceiving the world via the body can be investigated by presenting exteroceptive stimuli at different intervals from the heartbeat. For example, tones presented at short delays from the heartbeat (~250ms) are typically perceived as synchronous, while those presented at longer delays (~550ms) are typically perceived as asynchronous with the heart. Furthermore, when participants listen to sequences of such synchronous or asynchronous sounds, we previously observed an HEP effect in the period between the heartbeat and the expected sound, potentially reflecting cardio-audio integrated expectations (Banellis & Cruse, 2020). In support of attentional modulation of integrated predictive mechanisms, we also observed a larger positivity to unexpectedly omitted sounds in a sequence of cardio-audio synchronous sounds *only* when participants were attending to their heartbeat.

However, in that same study we found no evidence of evoked potential modulation by participants’ trait heartbeat perception abilities, thus failing to support the hypothesised trait precision contribution to HEPs (Katkin et al., 1991; Pollatos et al., 2005; Pollatos & Schandry, 2004; Schandry et al., 1986). One potential cause of this lack of evidence is that we failed to account for individual differences in the temporal location of heartbeat sensations (Brener & Ring, 2016). Multi-interval tasks, such as the Method of Constant Stimuli (MCS), are more sensitive at determining interoceptive ability as the optimal relative timing of heartbeat sensations is not presumed (Brener et al., 1993; Brener & Ring, 2016). For example, higher accuracy in a two-interval task can be achieved by first determining the optimal timing of each individual’s heartbeat sensations in a multi-interval task (Brener & Kluvitse., 1988a/b; Mesas & Chica., 2003).

Therefore, in this study, we sought to more sensitively test for trait precision influences on HEPs using the above method of individually-adjusted timings. Furthermore, we sought to replicate our previously observed effects of attention and cross-modal expectations on HEPs. Together, this study tests the hypotheses of predictive coding theories that HEPs reflect precision-weighted predictive mechanisms, where precision can be defined as both attentional gain and trait ability.

## Materials and Methods

Unless otherwise stated, all methods, analyses, and hypotheses were pre-registered at [https://osf.io/ptbzf/]

### Participants

Forty participants were recruited from the University of Birmingham via advertisement on posters or the online SONA Research Participation Scheme. Our inclusion criteria included: right-handed 18 to 35-year olds, with no reported cardiovascular or neurological disorders. We compensated participants with course credit or payment at a rate of £10 an hour. The STEM Research Ethics Board of the University of Birmingham granted ethical approval for this study and written informed consent was completed by all participants. Data of six participants were excluded due to EEG recording difficulties or poor data quality, resulting in more than a third of trials of interest rejected. One participant was rejected from part one (due to insufficient trials) but included in part two of the EEG analysis (and the opposite for a different participant). A final sample of 34 participants were included for both parts of subsequent EEG analyses (Median age = 20 years, Range = 18-35 years). This sample size was chosen in advance, as it provides 95% power to detect the same effect size (Cohen’s d’ = 0.58) as the within-subjects interaction between attention and cardio-audio delay observed in our previous experiment (preregistered analysis: M=0.0349, SD=0.0598; alpha=.05; note that the effect size in the final published version of that study [Banellis & Cruse., 2020] was slightly larger [0.61] due to pre-processing changes suggested by peer reviewers; GPower, (Faul et al., 2007)).

### Stimuli and Procedure

#### Overview

The experiment consisted of two parts; the function of part one was to determine the temporal location of perceived heartbeat sensations for each individual, using the Method of Constant Stimuli (MCS) (Brener et al., 1993; Brener & Kluvitse, 1988; Ring & Brener, 2018; Schneider et al., 1998). Part two comprised of a variant of a two-interval forced choice heartbeat discrimination task, with individually adjusted perceived synchronous and perceived asynchronous cardio-audio delays calculated from the median of their linearly interpolated cumulative distribution of choices from the MCS task (Brener & Kluvitse, 1988a, Brener & Kluvitse, 1988b; Mesas & Chica, 2003). Additionally, part two included an attention manipulation and interoceptive ability measurements, allowing the investigation of the effects of precision on cross-modal predictive mechanisms.

#### Part one: Method of constant stimuli

Part one consisted of three blocks of 40 trials (120 trials total), with each trial consisting of 5 to 7 auditory tones (1000Hz, 100ms duration, 44100 sampling rate) presented via external speakers, with breaks given between each block. The onset of each tone was triggered by the online detection of the participants R-peak from electrocardiography (ECG) recordings using Lab Streaming Layer and a custom MATLAB script (Kothe et al., 2018). The script analysed in real time the raw ECG signal by computing the variance over the preceding 33ms window and determining if the signal exceeded an individually adjusted threshold, at which point a tone was triggered to occur after one of six cardio-audio delays (an average time of 113ms, 213ms, 314ms, 413ms, 510ms, or 612ms delay). Due to computational variability in online detection of R-peaks, R->Sound intervals had a standard deviation of between 24ms-26ms. A fixation cross was present during tone presentation.

Participants focused on their heartbeat (without taking their pulse) and determined whether the tones presented were synchronous or not with their heartbeat. At the end of each trial, participants responded to the question ‘Were the tones synchronous with your heart?’ by pressing ‘y’ for yes or ‘n’ for no on the keyboard. The inter-trial interval was between 2 to 3 seconds, chosen from a uniform distribution on each trial (see Figure 1). The order of the experimental conditions was randomized to ensure no more than 3 of the same condition on consecutive trials.

**Figure 1.**
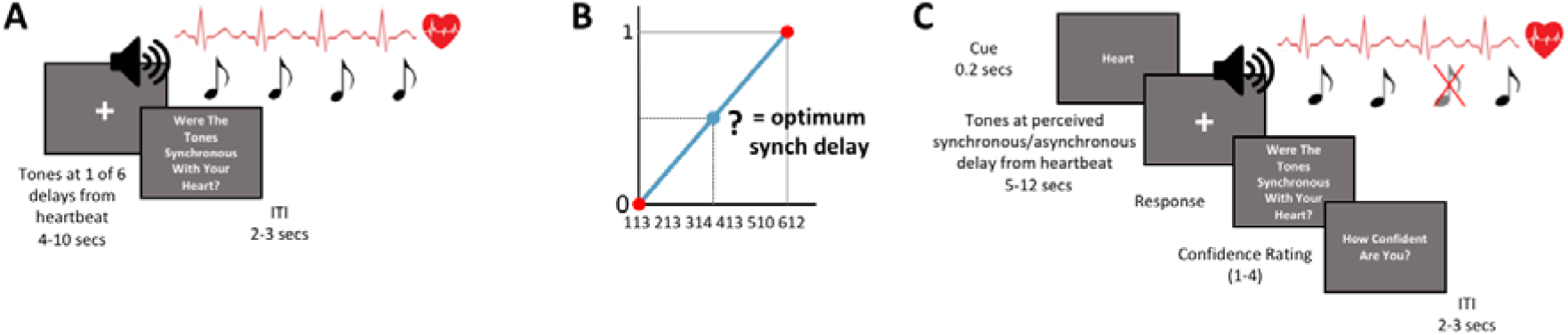
(A) Part 1 of the experiment consisted of a multi-interval heartbeat discrimination task (MCS) in which sounds were presented at 1 of 6 intervals from the heartbeat (113ms, 213ms, 314ms, 413ms, 510ms, or 612ms). (B) Calculation of each individuals perceived synchronous delay from the median of the linearly interpolated cumulative distribution of choices from the MCS task (marked as the question mark). (C) Part 2 of the experiment consisted of an individually adjusted two-interval heartbeat discrimination task for half of the trials (internal attention; as shown in C), and half consisted of an omission-detection task (external attention).

#### Part two: Individually-adjusted two-interval task

Part two consisted of three blocks of 56 trials (168 trials total), with each trial consisting of 7 to 10 auditory tones triggered by the online detection of the participant’s R-peak (as above in part one), presented via external speakers, with breaks given between each block. We selected each individual’s perceived synchronous cardio-audio delay from their performance in part one (i.e. the MCS), by calculating the median from the linearly interpolated cumulative percentage sum of their response counts for each delay (Brener & Kluvitse, 1988a, Brener & Kluvitse, 1988b; Mesas & Chica, 2003), and we selected the perceived asynchronous delay to be 300ms later (SD= 0.001) than the perceived synchronous delay, as in Mesas and Chica (2003). Due to computational variability in the online detection of R-peaks, R->Sound intervals had a standard deviation of 45ms for both the perceived synchronous and asynchronous trials. In half of the trials, we omitted the third from last tone, resulting in an R-peak without an auditory stimulus - an omission. We presented a fixation cross during tone presentation.

A cue at the start of each trial (200ms) directed participants’ attention to focus internally (‘Heart’) or externally (‘Tone’). During the internal task, participants focused on their heartbeat sensations (without taking their pulse) and determined whether the tones presented were synchronous or not with their heartbeat. During the external task, participants were told to ignore their heartbeat sensations and direct attention towards the sounds alone. The external task was to determine whether there was a missing sound during that trial. After each task response, participants rated their confidence in their decision from 1 to 4 (1 = Total Guess, 2 = Somewhat Confident, 3 = Fairly Confident, 4 = Complete Confidence). The inter-trial interval was between 2 to 3 seconds, chosen from a uniform distribution on each trial (see Figure 1). We randomized the order of the experimental conditions to ensure no more than 3 of the same condition on consecutive trials. Finally, participants completed the short Porges Body Perception Questionnaire (BPQ), including a body awareness and autonomic reactivity subscale (Porges, 1993).

### EEG/ECG acquisition

We recorded EEG throughout the experiment using a gel-based 128-channel Biosemi ActiveTwo system, acquired at 512Hz, referenced to the Common Mode Sense electrode located approximately 2-cm to the left of CPz. Two additional electrodes recorded data from the mastoids, and ECG was measured using two electrodes placed on either side of the chest, also sampled at 512Hz.

### EEG/ECG Pre-Processing

First, we filtered the continuous EEG data in two steps (i.e. low-pass then high pass) between 0.5Hz and 40Hz using the finite impulse response filter implemented in EEGLAB (function: pop_eegfiltnew). We filtered ECG between 0.5Hz and 150Hz (Kligfield et al., 2007) and in addition to that preregistered, we notch-filtered the ECG between 48Hz and 52Hz to remove line noise. Next, we segmented the filtered EEG signals into epochs from −300ms to 900ms relative to the R-peak of the ECG recording during within-task omission periods. In addition to that preregistered, we segmented EEG data during silent periods at the end of trials without an omission as equivalent to a within-task omission, to increase power. Endtrial silences are comparable to within-task omissions because participants could not predict when the trial would end due to the variable number of sounds in each trial. We segmented auditory-evoked potentials (AEPs) from −500ms to 500ms relative to the sounds during the MCS task and segmented HEPs −300 to 900ms relative to the first R-peak during end trial silent periods of the MCS task.

Initially, we re-referenced EEG data to the average of the mastoids. We detected the R-peaks using a custom MATLAB script, and subsequently checked the accuracy of R-peak detection via visual inspection. When necessary, we manually corrected the estimated timing of R-peaks to ensure accurate R-peak detection. To account for online heartbeat detection errors (i.e. missed or multiple sounds per R-peak), we rejected blocks with R-R intervals > 1.5 seconds or < 0.4 seconds from both behavioural and EEG analyses. In addition to that preregistered, to avoid contaminating responses within the analysis window, we rejected trials with triggers within 100ms prior to ERP onset (i.e., contaminating sounds for HEP trials and heartbeats for AEP trials). For AEPs this included the rejection of trials with R-peaks within the analysis window (sound-500ms). The subsequent artefact rejection proceeded in the following steps based on a combination of methods described by Mognon et al., 2011 and Nolan et al., 2010.

First, bad channels were identified and removed from the data. We consider a channel to be bad if its absolute z-score across channels exceeds 3 on any of the following metrics: 1) variance of the EEG signal across all time-points, 2) mean of the correlations between the channel in question and all other channels, and 3) the Hurst exponent of the EEG signal (estimated with the discrete second order derivative from the Matlab function *wfbmesti*). After removal of bad channels, we identified and removed trials containing non-stationary artefacts. Specifically, we considered a trial to be bad if its absolute z-score across trials exceeds 3 on any of the following metrics: 1) the mean across channels of the voltage range within the trial, 2) the mean across channels of the variance of the voltages within the trial, and 3) the mean across channels of the difference between the mean voltage at that channel in the trial in question and the mean voltage at that channel across all trials. After removal of these individual trials, we conducted an additional check for bad channels, and removed them, by interrogating the average of the channels across all trials (i.e. the evoked response potential (ERP), averaged across all conditions). Specifically, we considered a channel to be bad in this step if its absolute z-score across channels exceeds 3 on any of the following metrics: 1) the variance of voltages across time within the ERP, 2) the median gradient of the signal across time within the ERP, and 3) the range of voltages across time within the ERP.

To remove stationary artefacts, such as blinks and eye-movements, the pruned EEG data is subjected to independent component analysis (ICA) with the *runica* function of EEGLAB. The Matlab toolbox ADJUST subsequently identified which components reflect artefacts on the basis of their exhibiting the stereotypical spatio-temporal patterns associated with blinks, eye-movements, and data discontinuities, and the contribution of these artefact components is then subtracted from the data (Mognon et al., 2011). Next, we interpolated the data of any previously removed channels via the spherical interpolation method of EEGLAB, and re-referenced the data to the average of the whole head.

We subjected the data to a second round of ICA, to remove the CFA. This deviated from our preregistration, as ICA was deemed a more stringent CFA correction method than subtracting a rest template (see Supplementary Figure 8) and is the most frequently used CFA-correction method in the HEP literature (Coll et al., 2021; Park & Blanke, 2019). First, for the ICA computation, we filtered the ECG between 0.5Hz and 40Hz to ensure equivalent filtering as the EEG data and segmented the data into smaller epochs (−200ms to 200ms) with respect to the R-peak. We completed ICA on the shorter epoched data using the runica function of fieldtrip. To prevent multiple components with identical or symmetrical topographies, we set the maximum number of components to the rank of the data after trials concatenated. To select CFA components, we computed the pairwise phase coherence (PPC) of each component with the ECG. We selected a component if its PPC exceeded three standard deviations above the mean PPC of all components within the 0-25Hz range. We completed this selection procedure iteratively until no more than three components were selected. After visual inspection to ensure non-neural components had been identified, we removed the selected components from the original −300ms-900ms pre-processed EEG data. Finally, we visually inspected the data before and after CFA-correction and if the CFA was not visually diminished, we completed the cardiac ICA procedure again with an increased maximum number of rejected components, up to a maximum of six. The median number of components rejected across participants was 3 (range = 1-6).

Before proceeding to group-level analyses, we finalised single-subject CFA-corrected averages for HEP analysis in the following way. First, we generated a robust average for each condition separately, using the default parameters of SPM12. Robust averaging iteratively down-weights outlier values by time-point to improve estimation of the mean across trials. As recommended by SPM12, we low-pass filtered the resulting HEP below 20Hz (again, with EEGLAB’s pop_neweegfilt). In deviation from our pre-registration, but following discussions with peer reviewers and investigation of similar decisions in previous studies of HEPs (Azzalini et al., 2019; Babo-Rebelo et al., 2016, 2019; Banellis & Cruse, 2020; Park et al., 2014; Petzschner et al., 2019), we chose not to apply any baseline correction to our data as cardiac activity is cyclical by nature and may therefore insert artefactual effects in post-R data. This decision also allows direct comparison with published results of Banellis and Cruse (2020).

Finally, we visually inspected averaged ERPs to ensure the automated artefact rejection procedure was successful. If excessively large voltages remained in the averaged ERP, we visually inspected individual trials to ensure that any remaining channels/trials with excessively large voltages were removed. Additionally, if oculomotor artefacts remained then we identified additional ICA components and removed them manually.

### ERP analysis

We compared ERPs with the cluster mass method of the open-source Matlab toolbox FieldTrip (Oostenveld et al., 2011: fieldtrip-20181023). This procedure involves an initial parametric step followed by a non-parametric control of multiple-comparisons (Maris & Oostenveld, 2007). Specifically, we conducted either two-tailed dependent samples t-tests (part 1 AEPs, part 1 HEPs, and part 2 attention and delay comparisons, see Supplementary Table 1: comparisons 1, 2, 3 and 4) or a combination of two-tailed independent and dependent samples t-tests (part 2 delay and interoceptive ability, and part 2 attention and interoceptive ability comparisons, see Supplementary Table 1: comparisons 5 and 6) at each spatio-temporal data-point within time window of interest. We clustered spatiotemporally adjacent t-values with p-values < 0.05 based on their proximity, with the requirement that a cluster must span more than one time-point and at least 4 neighbouring electrodes, with an electrode’s neighbourhood containing all electrodes within a distance of .15 within the Fieldtrip layout coordinates (median number of neighbours = 11, Range = 2-16). Finally, we summed the t-values at each spatio-temporal point within each cluster. Next, we estimated the probability under the null hypothesis of observing cluster sum Ts more extreme than those in the experimental data - i.e. the p-value of each cluster. Specifically, Fieldtrip randomly shuffles the trial labels between conditions, performs the above spatio-temporal clustering procedure, and retains the largest cluster sum T. Consequently, the p-value of each cluster observed in the data is the proportion of the largest clusters observed across 1000 such randomisations that contain larger cluster sum T’s.

For the HEP analyses, to account for the lag difference in tone presentation across delay conditions, we completed one set of analyses on HEP data before omission onset relative to the R-peak (part 1: R+0-113ms post-R (i.e. earliest perceived synchronous cardio-audio delay); part 2: 0-129ms post-R (i.e. first percentile of the R->sound intervals for the participant with the earliest perceived synchronous delay, thus before >99% of anticipated tones) and a second set of analyses relative to the onset of the omitted sound (part 1: 0-250ms relative to the most rated and least rated synchronous delay time; part 2: 95-138ms post-omission (i.e. significant attention and delay interaction from Banellis & Cruse., 2020)). This allows the investigation of cardio-audio expectation and unfulfilled expectation mechanisms separately. The AEP analysis windows were determined by the global field power (GFP) and global mass dissimilarity (GMD) of the most and least rated synchronous conditions together (i.e. first three components preregistered: 0-74ms, 74-154ms, 154-209ms, fourth and fifth component exploratory: 209-289ms, 289-500ms). See Supplementary Table 1 for analysis details for all comparisons.

### Control Analyses

We performed a myriad of control analyses. This included analyses on physiological data (ECG, interbeat intervals (IBI’s), heart rate variability (HRV)), as well as additional analyses on EEG data (HEP control analyses and analytical controls such as baseline correction and CFA correction control analyses). For details of all control analyses and control results, see Supplementary Material.

### Interoceptive ability correlations

We correlated interoceptive ability (awareness, accuracy) with the mean difference in voltage across cardio-audio delay conditions during internal trials, using the electrodes and time-windows that demonstrated a significant main effect of cardio-audio delay (79-128ms post-R, and 94-137ms post-omission). In addition to that preregistered, to explore all aspects of interoceptive ability, we applied the same correlations during external trials and assessed the relationship of the above effects with interoceptive sensibility (body awareness and autonomic reactivity BPQ subsection separately), resulting in a total of 16 correlations.

### Source Reconstruction

In addition to the analyses preregistered, we performed source reconstruction to estimate the neural origin of each significant ERP effect using the open-source software Brainstorm, which implements a distributed dipoles fitting approach (Tadel et al., 2011). We completed our source estimation approach for each time-window separately in which we observed a significant sensor level effect. Specifically, within Brainstorm, we computed a forward model using the Symmetric Boundary Element Method (BEM) as implemented in OpenMEEG, based on the default MRI anatomy (ICBM152). We imported into Brainstorm each participant’s pre-processed EEG data prior to robust averaging, grouped by each attention and delay pair condition. We used a standard 128 Biosemi electrode position file for all participants. We generated the inverse model based on a minimum-norm solution using the current density map measure and unconstrained orientations, with an equal noise covariance matrix. We computed a grand average of the source results for each condition and subsequently averaged across the time window of each significant sensor-level effect. We calculated the difference between source maps and subsequently computed the norm of the three orientations, thus reflecting changes in source amplitude and orientation but not sign between source maps. We projected the estimated source results onto a canonical inflated brain surface for visualisation (plotting parameters: local maximum, amplitude=70%, minimum cluster=5).

## Results

### Behavioural data

#### Part one: MCS performance

We calculated the percentage preference for each delay by dividing the simultaneous judgement counts (total ‘yes’ responses for each delay trial) by the total trials for each delay after the removal of faulty blocks (R-R intervals > 1.5 seconds or < 0.4 seconds, as described above). Across participants, the mean delay at which sounds were perceived synchronous with the heart was 295.686 (SD=39.081). We classified a good heartbeat perceiver in part one on the basis of a Chi^2^ test which determined if the distribution of each individual’s simultaneous judgements deviated significantly from chance (Ring & Brener, 2018). This revealed a significant Chi^2^ effect for 8/34 participants for part 1 and part 2 each, and therefore defined 9 high heartbeat perceivers in total and 26 low heartbeat perceivers (see Supplementary Table 2 for individual performance).

#### Part two: Internal performance

Interoceptive accuracy (internal task d-prime), calculated from hits (responding ‘yes’ to a short ‘perceived synchronous’ cardio-audio delay trial) and false alarms (responding ‘yes’ to a long ‘perceived asynchronous’ cardio-audio delay trial; Macmillan & Creelman, 1990), was significantly greater than zero on average across the group (Mdn=0.271, Range=-0.445-2.166; Z = 530, *p* < .001, r_rb_ = 0.782), indicating that the sounds presented at individually adjusted cardio-audio delays were successfully perceived as synchronous and asynchronous. However, a Bayesian equivalent analysis indicated the evidence was weak relative to the null (BF_10_ = 2.876).

To determine whether individually-adjusted cardio-audio delays improved heartbeat perception, we compared internal performance here with that reported in an equivalent previous experiment without individually-adjusted delays (Banellis & Cruse, 2020). While interoceptive accuracy was more variable and had a higher median in this experiment (Mdn=0.271, Range=-0.445-2.166), than in the previous experiment with fixed delays (Mdn=0.204, Range=-0.447-1.274), this difference was not statistically significant (U = 502, *p* .134, r_rb_ = −0.156, BF_10_ = 0.463; note this remains non-significant with the removal of outliers).

#### Part two: External performance

Exteroceptive accuracy (external task d-prime), calculated from hits (responding ‘yes’ to a trial including an omission) and false alarms (responding ‘yes’ to a trial without an omission), was significantly greater than zero on average across the group (Mdn=3.134, Range=1.334-4.520; Z = 630, *p* < .001, r_rb_ = 1.000, BF_10_ = 93.999), demonstrating that participants were attentive, as required by the task demands.

There was no significant difference between the external accuracy scores during perceived synchronous trials (M=3.011, SD=0.804) and perceived asynchronous trials (M=2.918, SD = 0.934; t(34) = 0.836, *p* = .409, Cohen’s d’ = 0.141, BF10 = 0.251), indicating that external performance was not influenced by heartbeat perception. There was no significant correlation between internal and external performance (r_s_ = 0.131, *p* = .455, BF_10_ = .291), further signifying that internal and external task performance is unrelated.

### Event-related potentials

#### Part one: Method of constant stimuli

To test our hypothesis of reduced prediction error for sounds perceived as synchronous with the heart, we compared AEPs during cardio-audio delay trials most rated as synchronous with delay trials least rated as synchronous (comparison 1; see Supplementary Table 1 for analysis details). However, contrary to our hypothesis, we observed no significant AEP differences across perceived synchrony conditions (smallest *p* = .190).

Because there is an implicit omission for the first heartbeat after the end of each MCS trial, we also compared HEPs during periods of silence after the presentation of sounds to further test the hypothesis that HEPs reflect cardio-audio expectations. Specifically, we expected the first HEP during silent periods following a stream of stimuli perceived to be in cardio-audio synchrony to be larger relative to the HEP following stimuli perceived as asynchronous with the heartbeat. However, cluster-based permutation tests failed to reveal evidence of such expectation effects in this analysis when comparing the most rated synchronous trials with the least rated (R-locked: no clusters; omission-locked *p* = .248). Nevertheless, as the majority of participants (26) displayed a distribution of simultaneous choices at chance, it may not be meaningful to compare the most and least rated synchronous trials in this way. Subsequently, we exploratorily compared the MCS interval closest to the median of their simultaneous judgements with a 300ms later perceived asynchronous interval (as in the Part two HEP comparisons). While we didn’t observe a pre-omission HEP effect of perceived synchrony in this analysis (small *p* = .155), we did observe an end-trial omission-locked perceived effect of synchrony (cluster extending 176-248ms, positive cluster *p* = .004), consistent with HEPs reflecting processes linked to cardio-audio integration.

As an exploratory analysis, we computed the above in high and low perceivers separately, as defined by the Chi^2^ of each individual’s MCS performance (comparison 3). When analysing AEPs in high heartbeat perceivers only (although only a small sample of 8 participants in part one, determined by a Chi^2^ on individual MCS performance), we observe a larger early fronto-central positivity for trials perceived as synchronous (cluster extending 176-209ms, positive cluster *p* = .021), followed by a larger fronto-central negativity for perceived asynchronous trials (cluster extending 240-289ms, positive cluster *p* = .007) consistent with cardiac-driven auditory prediction error. As all equivalent comparisons in low perceivers were not significant (smallest *p* = .311), this result is also consistent with a role of trait precision on HEP amplitude.

Source estimates of the initial AEP effect demonstrated the largest clusters in the left inferior frontal cortex and left temporopolar area, with smaller clusters including left premotor cortex and left primary sensory cortex. Source analysis of the following AEP effect demonstrated the largest clusters in the left inferior frontal cortex, bilateral anterior frontal cortex and left temporopolar area, with smaller clusters including left premotor cortex, right primary sensory cortex, bilateral visual association area.

All other ERP comparisons for part one (MCS) high perceivers were not significant (smallest *p* = .046).

#### Part two: Individually-adjusted two-interval task

##### Cardio-audio expectation

To test our hypothesis of cardiac-driven expectations of sounds, we compared HEPs across cardio-audio delay conditions pre-omission. We observed a pre-omission main effect of delay (positive cluster *p* = .024) locked to the R-peak, replicating our previous finding with fixed cardio-audio delays and supporting our hypothesis of heartbeat-driven predictions of auditory stimuli (Banellis & Cruse, 2020). The positive cluster extended from 79-128ms. Source estimates of this effect include largest clusters in left middle temporal gyrus and right supramarginal gyrus and smaller clusters in bilateral frontal eye fields, left dorsolateral prefrontal cortex, left visual association area, right superior temporal gyrus and right fusiform gyrus.

To test our hypothesis of attentional precision modulating predictive mechanisms, we calculated the attention and delay interaction as the difference between short-delay and long-delay trials between attention groups (i.e. a double-subtraction; comparison 4). However, we did not observe a significant R-locked delay and attention interaction (*p* = .401). Nevertheless, we observed a significant main effect of attention on pre-omission responses (negative cluster *p* = .013, cluster extending 37-68ms). Source estimates of this effect include largest clusters in left anterior prefrontal cortex and right visual association area, with smaller clusters in left dorsolateral prefrontal cortex and right fusiform gyrus.

### Unfulfilled expectation

Inconsistent with our hypothesis of attentional precision modulating predictive mechanisms, and inconsistent with evidence from a previous study (Banellis and Cruse, 2020), the attention and delay interaction for omission-locked responses was also not significant (*p* = .159). One potential cause of this lack of replication is that in this experiment we defined omissions to include not only within-task silent periods, but also silent periods at the end of trials without an omission to increase power. However, when we selected within-task omissions only and analysed the significant electrodes and time window of the delay and attention interaction from our previous study (Banellis & Cruse, 2020) this interaction is also not significant (F(1,32) = 2.141, *p* = .153, n^2^ = 0.022, BF_incl_ = 0.100).

To test our hypothesis of higher prediction error during omission periods in a stream of auditory stimuli perceived as synchronous with the heart, we compared omission-evoked responses across cardio-audio delay conditions. We observed an omission-locked main effect of delay, with the positive cluster extending 94-137ms (positive cluster *p* = .022). Source estimates include largest clusters in left inferior frontal gyrus, right anterior frontal cortex, with smaller clusters in left superior temporal gyrus.

### Interoceptive ability

For all evoked potential analyses, we separated our participants into groups of high/low interoceptive accuracy, sensibility, and awareness with median splits. As mentioned, we defined *interoceptive accuracy* as the difference between the normalised proportion of hits and the normalised proportion of false alarms (i.e. internal task *d*’, see above) (Macmillan & Creelman, 1990). As in previous studies (Ewing et al., 2017; Garfinkel et al., 2015), we quantified sensibility to a variety of internal bodily sensations with the score on the awareness subsection of the Porges Body Perception Questionnaire (BPQ) (Porges, 1993) and defined *sensibility* to heartbeat sensations as the median confidence rating during internal trials (Ewing et al., 2017; Forkmann et al., 2016; Garfinkel et al., 2015). We also calculated *interoceptive awareness* using type 2 signal detection theory analysis comparing observed type 2 sensitivity (meta-*d*’) with expected type 2 sensitivity (*d*’) (i.e. meta-d’ - d’) (Maniscalco & Lau, 2012). Meta-*d*’ is the *d*’ expected to generate the observed type 2 hit rates and type 2 false alarm rates and was estimated using maximum likelihood estimation (MLE) (Maniscalco & Lau, 2014). This determined the extent to which confidence ratings predicted heartbeat detection accuracy, and thus interoceptive awareness.

First, we tested our hypothesis of interoceptive ability modulating the attention effect observed previously in Banellis & Cruse (2020) (comparison 6). We observed a significant omission-locked interaction of interoceptive awareness and attention during synchronous trials (positive cluster *p* = .014, cluster extending 96-139ms). Source estimates of this effect include right frontal eye fields and bilateral visual association cortex. Pairwise comparisons of omission responses during synchronous trials revealed a significant difference between attention conditions in participants with high interoceptive awareness (negative cluster *p* = .019, cluster extending 105-131ms); no clusters were observed when comparing low awareness participants. Source estimates of the attention effect in high awareness participants reveal the left anterior frontal cortex, left dorsolateral prefrontal cortex and right visual association cortex. Source estimates of the same time-window in the low awareness group includes bilateral visual association cortex, right angular gyrus and right fusiform gyrus.

All other omission-locked interoceptive ability interactions with attention during synchronous trials were not significant (interoceptive accuracy (smallest *p* = .097), interoceptive sensibility: median confidence (smallest *p* = .161), the awareness subsection (smallest *p* = .081) and the autonomic reactivity subsection (smallest *p* = .061) of the BPQ separately). Additionally, no significant R-locked interoceptive ability and attention interactions during synchronous trials were observed (smallest *p* = .099).

Next, we tested our hypothesis of interoceptive ability modulating the delay effect (comparison 5) and observed no omission-locked interactions during internal trials (interoceptive accuracy (no clusters), awareness (no clusters) or sensibility (median confidence (smallest *p* = .127), the awareness (smallest *p* = .350) and the autonomic reactivity (smallest *p* = .210) subsection of the BPQ separately). Additionally, no significant R-locked interoceptive ability and delay interactions during internal trials were observed (smallest *p* = .107).

Finally, we observed no significant correlations of interoceptive ability with the amplitude of the omission-locked delay effect (smallest *p* = .184). However, we observed an uncorrected significant correlation of the awareness subsection of the BPQ and the R-locked delay effect during external attention (*p* = .022), however this is 1 out of the 16 correlations (bonferroni corrected alpha = .003). All other correlations of interoceptive ability and the R-locked delay effect were not significant (smallest *p* = .233).

## Discussion

Interoceptive and exteroceptive integration is fundamental for the interwoven interactive experience of the body with the external world. These integrated signals are proposed to operate predictively, with regulation by precision-weighting (Barrett & Simmons, 2015; Cameron, 2002; Seth, 2013; Seth et al., 2012; Seth & Friston, 2016). In a previous study, we observed integrated cardio-audio predictive mechanisms by studying HEPs during heartbeat-predicted omissions (Banellis & Cruse., 2020). While our data in that study were consistent with the modulation of HEPs by attentional precision, we found no evidence of the influence of trait precision - i.e., individual interoceptive ability - contrary to the expectations of predictive coding. Consequently, in this study, we tailored the cardio-audio delays used for each individual to more accurately investigate trait-precision modulations of predictive signals, and subsequently determine if intero-extero integration operates in accordance with the predictive coding framework.

Despite our use of an arguably more sensitive and individually-tailored heartbeat perception task, we found no evidence for an HEP relationship between any measure of interoceptive ability and cardio-audio delay. One interpretation is that this may be due to the difficulties of assessing interoceptive performance, as we assess this indirectly with a relatively difficult task. For example, even with a more sensitive measure of objective performance across multiple cardio-audio delay intervals, only 9/35 participants were classified as high heartbeat perceivers. Additionally, influences of interoceptive ability may occur much later than can be observed with our design. For example, ERPs associated with metacognition often occur up to 1900ms post-stimulus, thus overlapping with forthcoming heartbeats and/or sounds (Skavhaug et al., 2010; Sommer et al., 1995; Tsalas et al., 2018). Furthermore, metacognitive awareness may be reflected in other features of the EEG, such as global long-range connectivity patterns, rather than local HEP differences (Canales-Johnson et al., 2015). Our specific HEP results here, nevertheless, fail to support a predictive coding account of interoceptive-exteroceptive integration under which predictive processes are modulated by trait-level precision.

Furthermore, we also failed to replicate the previously reported attention and delay interaction of omission-evoked potentials, contrary to a predictive coding account in which attention modulates expectations by precision-weighting. One possible interpretation is that, in this study, participants relied less on attentional-precision to ‘boost’ their predictions due to the enhanced perception of cardio-audio synchrony, reflected in the trend for increased performance relative to the previous experiment (see Figure 2C). As a result, attentional modulations of HEPs may have been weaker in this study. Despite this, we did observe a significant omission-locked delay effect, demonstrating the presence of cardio-audio predictive mechanisms, although without evidence of attentional modulation (see Figure 6). This is comparable to findings by Pfieffer and De Lucia (2017) who also found an HEP difference during omission periods when comparing cardio-audio synchronous streams with asynchronous streams in participants who were not actively attending to cardio-audio synchronicity. However, in that study, due to the timing of the auditory stimuli, it was not possible to separate omission-evoked effects from expectation effects. While we overcame this in our study by employing cardio-audio delays, allowing for the independent investigation of expectation and unfulfilled expectation effects, we also observe no evidence of the necessity of attention for generating auditory expectations on the basis of the heartbeat. Indeed, despite our previous observations (Banellis and Cruse, 2020), our Bayesian analysis in this study indicated strong evidence (i.e. BF=10 in favour of the null) for the absence of an interaction with attention - inconsistent with a predictive coding account.

**Figure 2.**
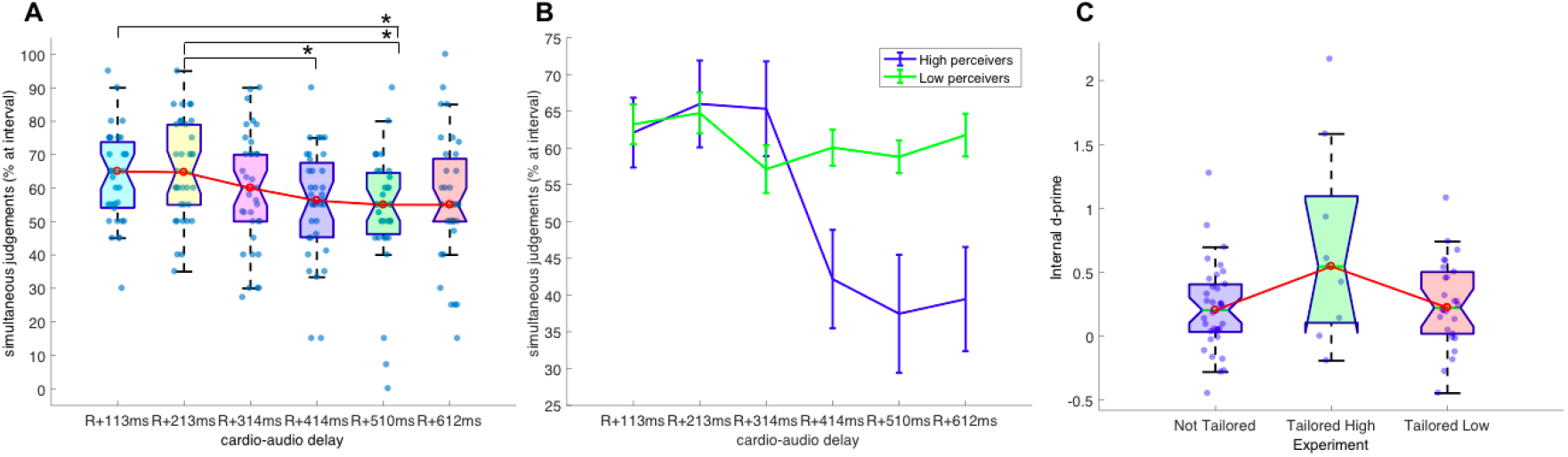
(A) boxplots of percentage preferences for each cardio-audio delay during the MCS, asterisks mark significant differences. (B) mean percentage preferences during the MCS split into high and low perceivers using Chi2, error bars reflect standard error of the mean. (C) boxplots of internal d’ performance comparing experiment 1 (‘Not Tailored’ delays: Banellis & Cruse., 2020) with this experiment (‘Tailored’ delays) split into high and low heartbeat perceivers using Chi2 of MCS performance.

**Figure 3.**
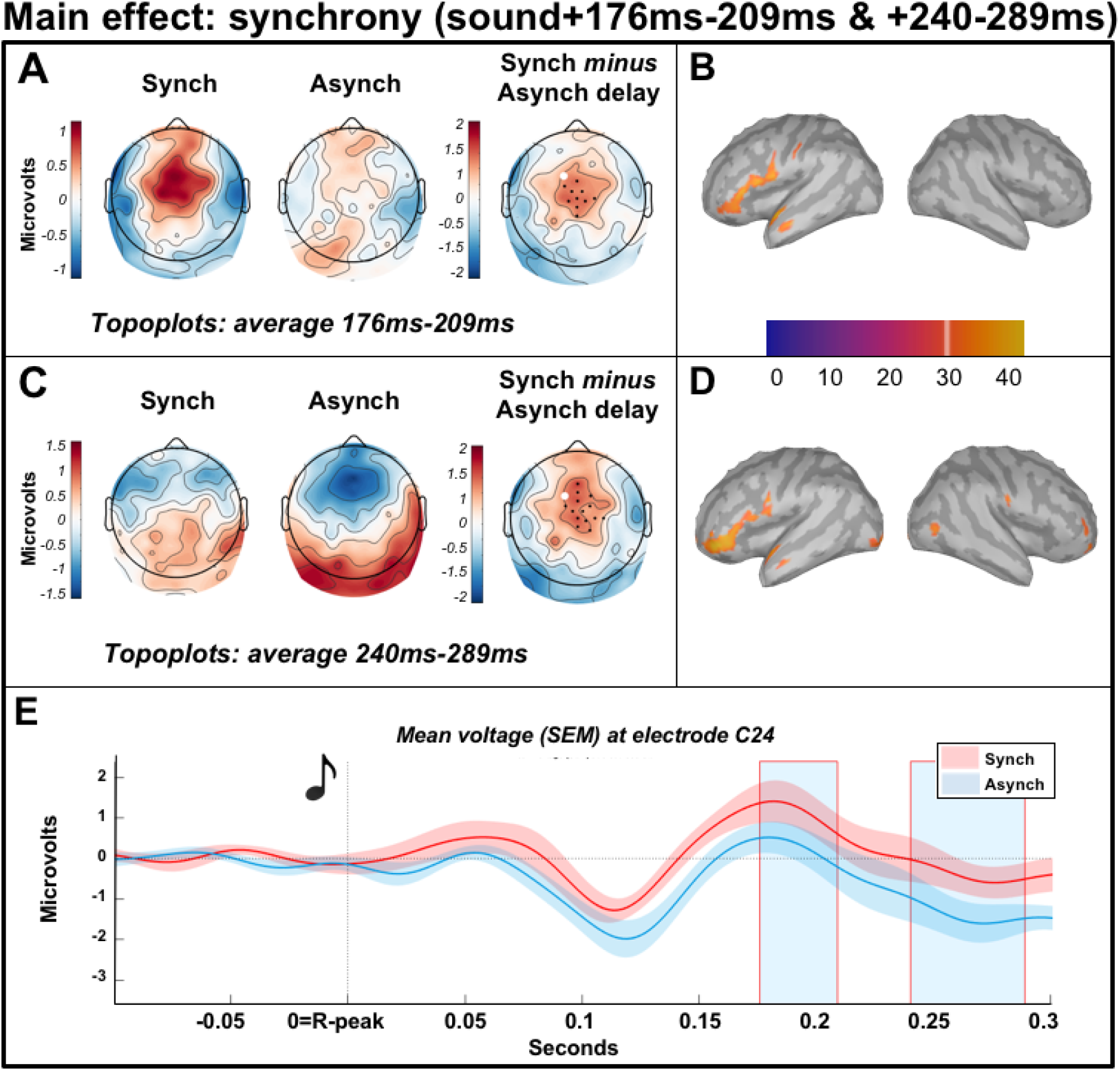
Main effect of perceived synchrony from 176 to 209ms and 240 to 289ms relative to most (synch) and least (asynch) rated synchronous sounds during the MCS task (part 1), in high heartbeat perceivers only. Scalp distribution of the average significant difference across perceived synchrony conditions (A) 176-209ms and (C) 240-289ms with electrodes contributing to the cluster marked. (B) Estimated sources of the 176-209ms main effect include largest clusters in the left inferior frontal cortex and left temporopolar area, smaller clusters included left premotor cortex and left primary sensory cortex. (D) Estimated sources of the 240-289ms main effect include largest clusters in the left inferior frontal cortex, bilateral anterior frontal cortex and left temporopolar area, smaller clusters included left premotor cortex, right primary sensory cortex, bilateral visual association area. (E) average AEP across participants at electrode C24, light blue shaded regions represent the time of the significant positive effect.

Upon visual inspection of our data, we were concerned about the presence of HEP differences prior to omissions in some comparisons, in particular that shown in Figure 6. These baseline differences may subsequently confound any apparent post-omission effects. Due to the cyclical nature of the heartbeat, and to be consistent with some previous literature (Azzalini et al., 2019; Babo-Rebelo et al., 2016, 2019; Banellis & Cruse, 2020; Park et al., 2014; Petzschner et al., 2019), we did not apply baseline correction in our preprocessing above. However, this choice is not ubiquitous in the HEP literature. Indeed, the issues for replication that are posed by the range of pre-processing / analysis / CFA correction methods employed across the field have recently been highlighted (Coll et al., 2021; Park & Blanke, 2019). Consequently, we re-analysed all effects reported here using an additional five sets of pre-processing pipelines (e.g., with baseline correction / without CFA correction, etc.; see Supplementary Table 3 for details) to identify the consistency of our observed effects (Botvinik-Nezer et al., 2020; Simonsohn et al., 2015; Steegen et al., 2016). We were reassured to find that the post-omission delay effect remains significant across all pre-processing pipelines, strengthening our interpretation that it reflects cross-modal integrative predictive processes, rather than analytical confounds (see Supplementary Figure 9).

Additionally, we replicated our previously observed pre-omission HEP difference across cardio-audio delay trials, likely reflecting a difference in cardio-audio expectation and supporting the hypothesis of interoceptive signals guiding expectations of exteroceptive stimuli (see Figure 4). However, the scalp topography and estimated sources of the pre-omission delay effect here are not entirely overlapping with those observed previously. For example, although source estimates from both studies revealed the middle temporal gyrus, supramarginal gyrus, and broad frontal regions, somatosensory and motor regions were also evident in Banellis and Cruse (2020), while visual and fusiform areas were evident in this study only. One possible reason for this disparity is that the previously reported expectation effect (Banellis and Cruse, 2020) extended to 230ms post-R, while the pre-omission window in this study was necessarily shorter (R+129ms) due to our use of individualised delays. Nevertheless, the topographical differences across experiments persist even when using a shorter time-window in our previous study. It may therefore be that our use of tailored delays in this study enhanced heartbeat-driven expectations in more participants, as supported by the trend for better objective performance, thus more accurately reflecting cross-modal expectations and subsequent predictive sources.

**Figure 4.**
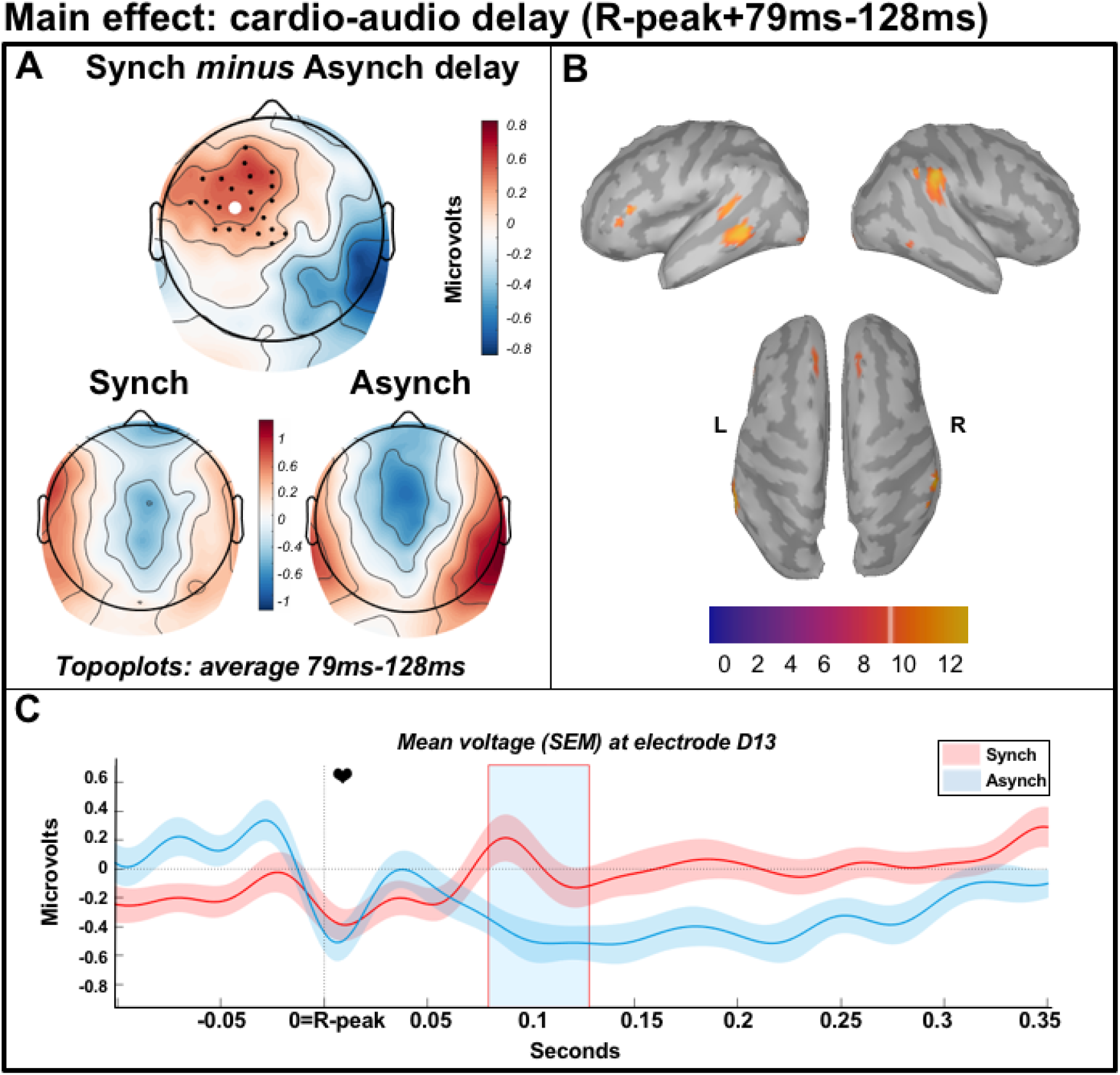
Main effect of cardio-audio delay from 79 to 128ms relative to R-peak during pre-omission periods in the individually-adjusted two interval task (part 2). (A) Scalp distribution of the average significant difference across delay conditions 79-128ms, with electrodes contributing to the cluster marked. (B) Estimated sources of the main effect include largest clusters in left middle temporal gyrus and right supramarginal gyrus and smaller clusters in bilateral frontal eye fields, left dorsolateral prefrontal cortex, left visual association area, right superior temporal gyrus and right fusiform gyrus. (C) average HEP across participants at electrode D13, light blue shaded region represents the time of the significant positive effect.

**Figure 5.**
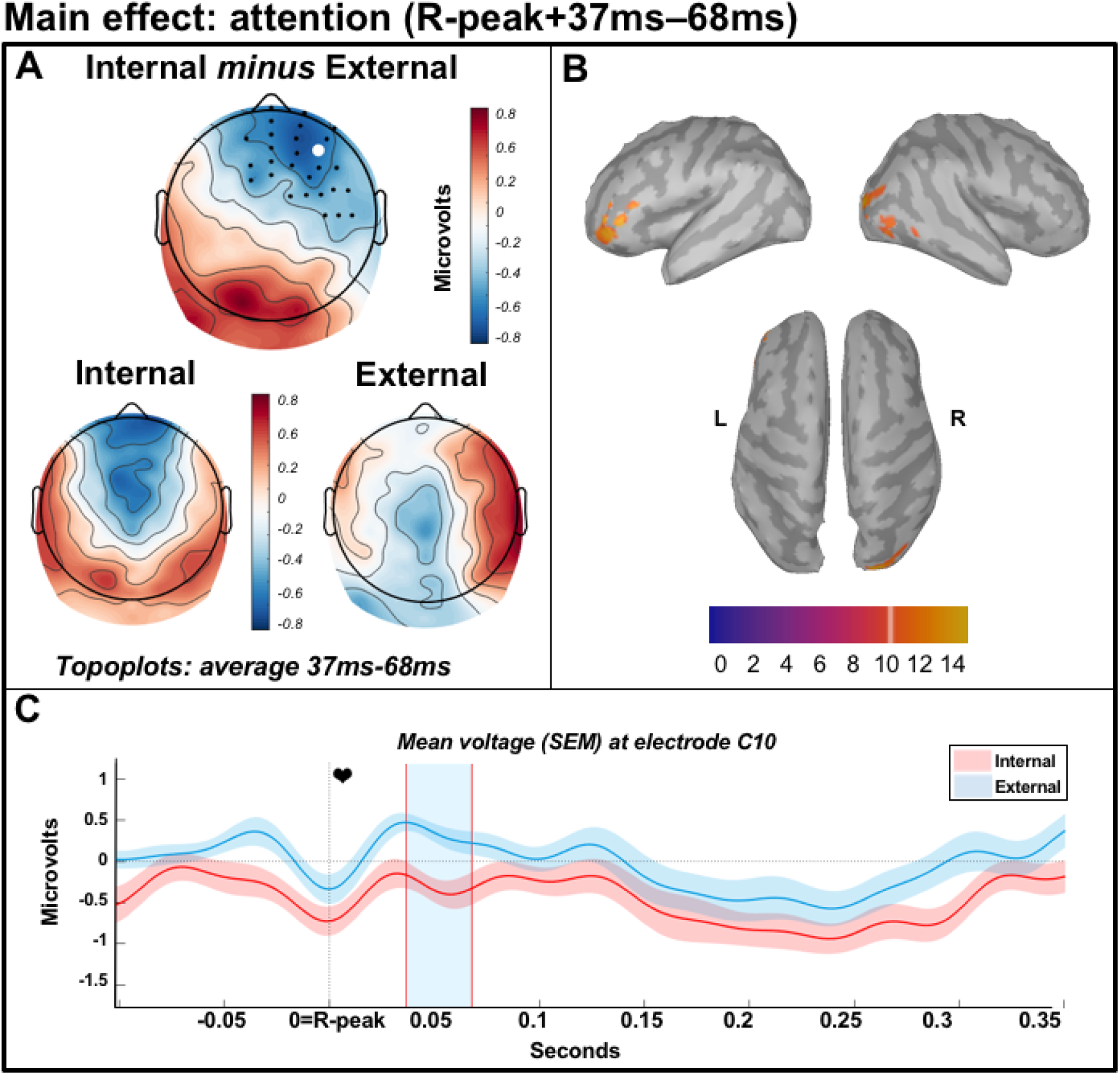
Main effect of attention from 37 to 68ms relative to R-peak during preomission periods in the individually-adjusted two interval task (part 2). (A) Scalp distribution of the average significant difference across attention conditions 37-68ms, with electrodes contributing to the cluster marked. (B) Estimated sources of the main effect include largest clusters in left anterior prefrontal cortex and right visual association area, smaller clusters in left dorsolateral prefrontal cortex and right fusiform gyrus. (C) average HEP across participants at electrode C10, light blue shaded region represents the time of the significant negative effect.

**Figure 6.**
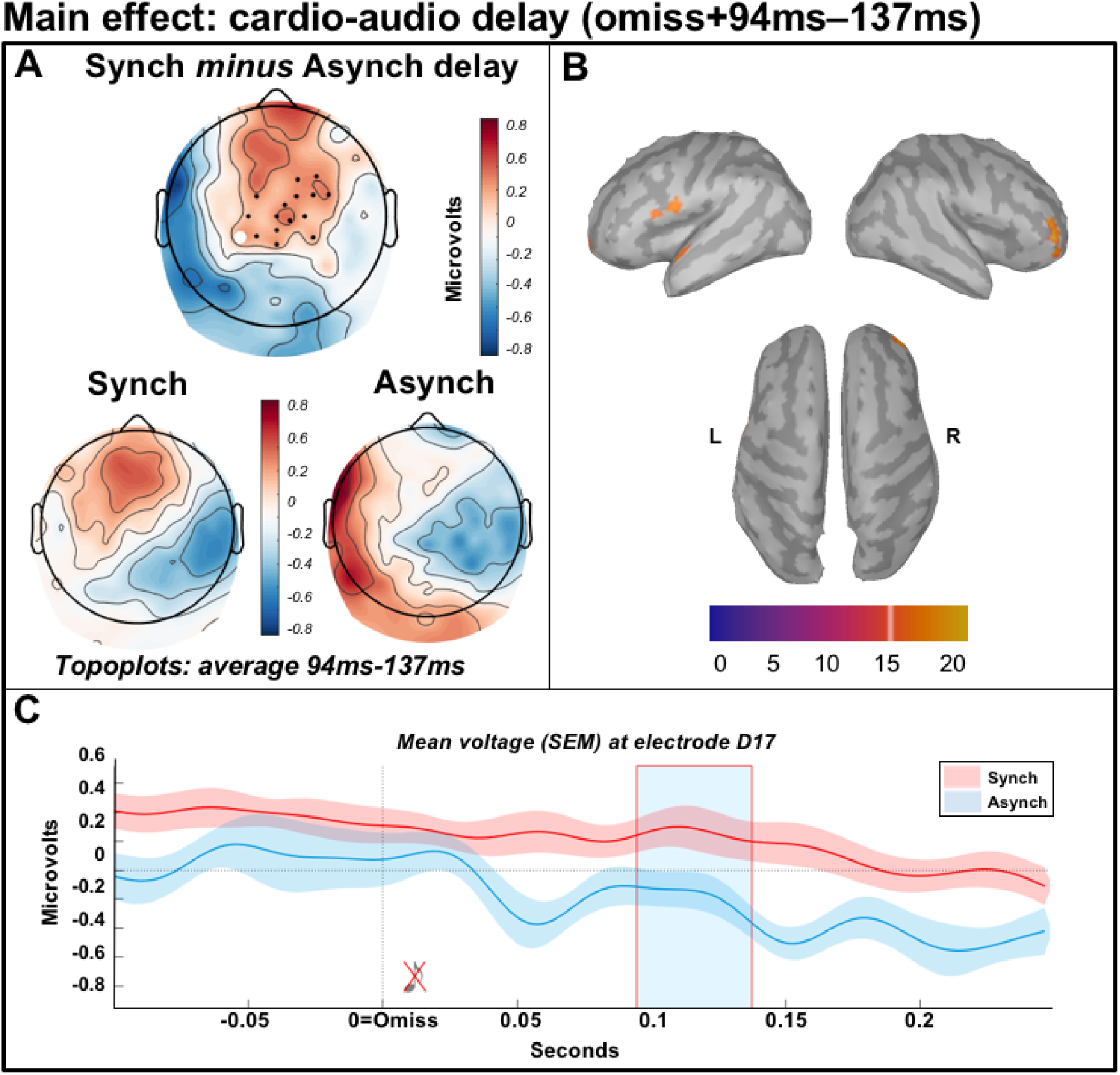
Main effect of cardio-audio delay from 94 to 137ms relative to the omission during the individually-adjusted two interval task (part 2). (A) Scalp distribution of the average significant difference across delay conditions 94-137ms, with electrodes contributing to the cluster marked. (B) Estimated sources of the main effect include largest clusters in left inferior frontal gyrus, right anterior frontal cortex, smaller clusters in left superior temporal gyrus. (C) average omission-evoked response across participants at electrode D17, light blue shaded region represents the time of the significant positive effect.

Although not interacting with cardio-audio delay, we did observe some evidence of the influence of interoceptive ability on HEPs in our omission-locked interaction of attention with interoceptive awareness (see Figure 7). This significant interaction reflected an attentional difference in high awareness participants only. Consistent with this result, previous research has reported a greater attentional HEP difference in good heartbeat perceivers, relative to poor perceivers (Montoya et al., 1993; Yuan et al., 2007). However, rather than the heartbeat discrimination task we employed here, those previous studies used the heartbeat counting task, which problematically confounds heartbeat perception with the ability to estimate heartrate or time (Brener & Ring, 2016; Ring & Brener, 2018). The effect observed here temporally overlaps with an effect of delay, potentially indicating that with high awareness, attention alters intero-extero predictive mechanisms. However, this effect was present in only a subset of the pre-processing pipelines, thus requiring cautious interpretation. Indeed, when studying neural activity time-locked to bodily events, it is crucial to test for the confounding influence of both peripheral physiological signals and analytical decisions. For example, we observed no heartrate or HRV differences in the directions of interest, and no ECG differences across conditions of interest for all analyses reported here, giving us confidence that our results reflect neural activity. Conversely, the behaviour of HEP effects across multiple pre-processing pipelines provides a valuable indicator of confidence in the observed effects. As described above, standardisation and understanding of HEP preprocessing and analyses are vital for the progress of the field (Bigdely-Shamlo et al., 2016; Coll et al., 2021; Farzan et al., 2017).

**Figure 7.**
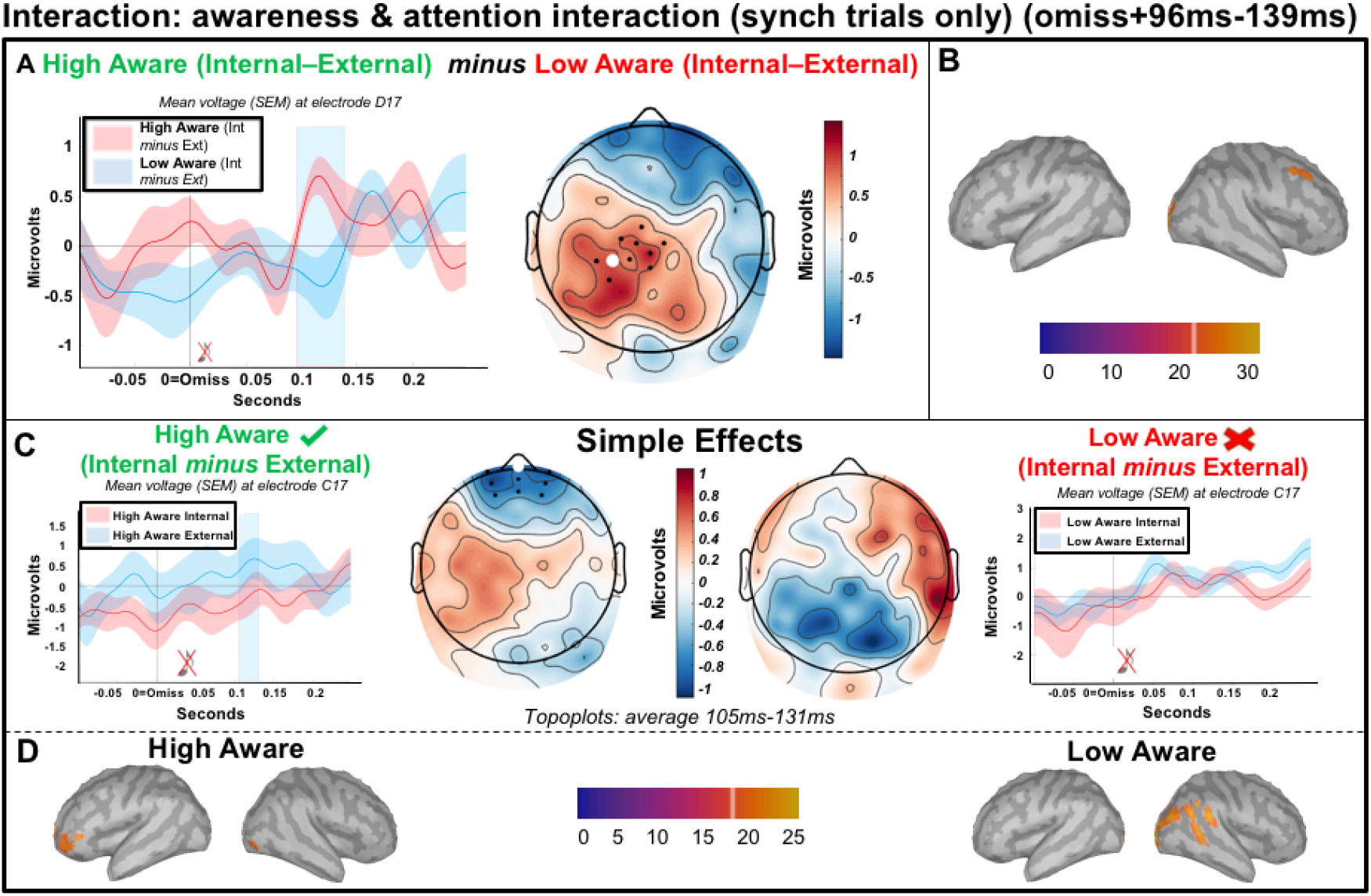
Interaction of interoceptive awareness and attention from 96 to 139ms relative to the omission during synchronous trials only in the individually-adjusted two interval task (part 2). (A - left) Average omission-evoked response across participants at electrode D17, light blue shaded region represents the time of the significant effect. (A-right) Scalp distribution of the average significant interaction (awareness x attention) 96-139ms, with electrodes contributing to the cluster marked. (B) Estimated sources of the interaction include right frontal eye fields and bilateral visual association cortex. (C) Analysis of the simple effects showing qualitatively different topographic distributions across interoceptive awareness groups (105-131ms) and a significant effect of attention in high awareness participants only. (D-left) Estimated sources of high awareness simple effects analysis reveal the left anterior frontal cortex, left dorsolateral prefrontal cortex and right visual association cortex. (D-right) Estimated sources of low awareness simple effects analysis includes bilateral visual association cortex, right angular gyrus and right fusiform gyrus.

Despite our lack of evidence for precision-weighting of HEPs by either attention or interoceptive ability, the robust pre- and post-omission delay effects observed here (and previously; Banellis and Cruse, 2020), are consistent with HEPs reflecting aspects of an integrated cardio-audio expectation process. Some accounts describe intero-extero expectation mechanisms as fundamental for embodied selfhood, emotion, and the generation of an integrated first-person perspective (Azzalini et al., 2019; Seth, 2013; Seth et al., 2012; Seth & Friston, 2016). Therefore, our paradigm may provide a tool for investigating cross-modal expectation processes in clinical conditions, as well as assessing its influence on cognition.

In conclusion, here we replicate evidence of cardiac signals guiding expectations of auditory stimuli. Despite this, we observe no evidence of either attentional-precision or traitprecision modulating these predictive processes, suggesting that intero-extero integration may not operate entirely within a precision-weighted predictive coding framework. Our results demonstrate a need for a clearer definition of the manipulation and measurement of precision on HEP effects, and the specific predictions made by predictive coding theories more generally. Finally, the robust delay effects observed here, and previously, may be useful for the investigation of the role of intero-extero integration in cognition, as well as for assessing its dysfunction in clinical groups.

## Author Notes

Funding: the Medical Research Council IMPACT Doctoral Training Programme at the University of Birmingham (scholarship to L.B.); Medical Research Council New Investigator Research (grant MR/P013228/1; to Principal investigator D.C.).

