## Supplementary material for "Heartbeat-evoked potentials during interoceptive-exteroceptive integration are not consistent with precision-weighting"

**Table 1.** *Details of ERP analyses.*

| **Comparison** | **Time window** | **Tail** |
| --- | --- | --- |
| 1. Perceived synchronous AEPs (most rated synchronous) and perceived asynchronous AEPs (least rated synchronous) during part 1. | First three auditory components determined by GFP and GMD: 0-74ms, 74-154ms, 154-209ms. Exploratory analysis of fourth and fifth auditory component: 209-289ms, 289-500ms. | Two-tailed. |
| 2. Perceived synchronous HEPs (most rated synchronous) and perceived asynchronous HEPs (least rated synchronous) during silent periods at the end of part 1 trials. | Separate time windows for the perceived synchronous cardio-audio delay (highest simultaneous judgement) and the perceived asynchronous cardio-audio delay (lowest simultaneous judgement) + 250ms each. For example, if the perceived synchronous delay was 213ms and the asynchronous delay was 510ms, the time windows would be 213-463ms and 510-760ms.  Also, R-peak to earliest perceived synchronous cardio-audio delay (113ms). | Two-tailed.  Two-tailed. |
| 3. Comparison 1 and 2 in high and low heartbeat perceivers separately (exploratory). | As comparison 1 and 2. | As comparison 1 and 2. |
| 4. Cardio-audio delay (perceived synchrony) and attention. 5. Interoceptive ability and cardio-audio delay, during internal trials. | R-peak to earliest perceived synchronous cardio-audio delay (129ms: defined as the 1^st^ percentile of the R->Sound intervals of p14 (participant with the lowest perceived synchronous delay).  Banellis and Cruse (2020) attention and cardio-audio delay interaction time window (95ms-138ms relative to the omission) | Two-tailed  One-tailed (perceived synchronous more positive) |
| 6. Interoceptive ability and attention, during synchronous trials. | R-peak to earliest perceived synchronous cardio-audio delay (129ms: defined as the 1^st^ percentile of the R->Sound intervals of p14 (participant with the lowest perceived synchronous delay).  Banellis and Cruse (2020) attention and cardio-audio delay interaction time window (95ms-138ms relative to the omission) | Two-tailed  Two-tailed |

**Table 2.** *Individual performance. ‘Med’ is the median (perceived synchronous) interval calculated from the linearly interpolated cumulative distribution of choices from the MCS task. ‘Max Int’ and ‘Min Int’ are the maximum and minimum delays rated as synchronous with the heartbeat during the MCS. ‘IQR’ is the difference between the 75^th^ and 25^th^ quartile of the linearly interpolated cumulative distribution of choices from the MCS task. To account for the* computational lag when triggering a sound from the online detection of R-peaks, *we added 113ms (i.e. average cardio-audio computational lag) to the Med and IQR columns, which were calculated from values inputted into the computer for the experiment. ‘Chi2’ on MCS simultaneous ratings: if significant (‘sig’) participants were classified as a high heartbeat perceiver, if not significant (‘not sig’) participants were defined as a low perceiver. ‘Internal d’’ (i.e. interoceptive accuracy) reflects d-prime of internal task performance. ‘External d’’ (i.e. exteroceptive accuracy) is the d-prime of external task performance. ‘BPQBA’ is the score on the Porges body perception questionnaire (short form) body awareness subsection, and ‘BPQANS’ is the score on the autonomic nervous system reactivity subsection (i.e. both reflect interoceptive sensibility to all body sensations). ‘Heart Conf’ is the median confidence rating during the internal task (i.e. interoceptive sensibility to heart sensations specifically). ‘Tone Conf’ is the median confidence ratings during the external task. ‘M_diff’ is meta d-prime, calculated as the difference between type 2 sensitivity (meta-d’) and expected type 2 sensitivity (d’) (i.e. meta-d’ – d’; interoceptive metacognitive awareness).*

| ID | Med | Max Int | Min Int | IQR | Chi2 | Internal d' | External d' | BPQBA | BPQANS | HeartConf | ToneConf | M_diff |
| --- | --- | --- | --- | --- | --- | --- | --- | --- | --- | --- | --- | --- |
| p04 | 309 | 612 | 510 | 423 | not sig | 0.589 | 2.848 | 69 | 37 | 2 | 4 | -0.666 |
| p05 | 248 | 213 | 612 | 362 | sig | 0.932 | 4.200 | 69 | 27 | 3 | 4 | -0.979 |
| p06 | 318 | 213 | 314 | 428 | not sig | 0.740 | 3.446 | 54 | 25 | 3 | 4 | -1.239 |
| p07 | 281 | 314 | 612 | 345 | sig | 0.000 | 2.590 | 47 | 24 | 2 | 3 | -0.087 |
| p08 | 299 | 213 | 612 | 399 | not sig | -0.123 | 1.334 | 98 | 36 | 3 | 3 | 0.653 |
| p09 | 301 | 113 | 314 | 428 | not sig | 0.131 | 3.118 | 83 | 39 | 2 | 3 | -0.246 |
| p11 | 325 | 213 | 113 | 432 | sig | 0.546 | 3.951 | 61 | 24 | 2 | 4 | -0.921 |
| p12 | 294 | 213 | 413 | 428 | not sig | 0.190 | 2.774 | 89 | 38 | 2 | 4 | -0.735 |
| p13 | 331 | 612 | 113 | 410 | not sig | -0.275 | 4.482 | 80 | 37 | 2 | 4 | 0.070 |
| p14 | 158 | 113 | 510 | 275 | sig | 2.166 | 2.433 | 69 | 39 | 4 | 4 | -0.903 |
| p15 | 294 | 113 | 314 | 469 | not sig | 0.272 | 1.570 | 30 | 23 | 2 | 3 | 0.181 |
| p17 | 329 | 113 | 314 | 454 | not sig | 0.019 | 3.425 | 53 | 33 | 2 | 4 | 0.278 |
| p18 | 354 | 510 | 213 | 415 | not sig | -0.184 | 1.589 | 84 | 35 | 3 | 3 | 0.175 |
| p19 | 301 | 113 | 612 | 422 | not sig | 0.672 | 2.724 | 67 | 28 | 2 | 3 | -0.557 |
| p20 | 309 | 213 | 612 | 385 | not sig | 0.456 | 2.517 | 94 | 31 | 3 | 3 | -1.210 |
| p21 | 271 | 314 | 510 | 344 | sig | 0.607 | 3.337 | 87 | 35 | 3 | 4 | -0.366 |
| p22 | 358 | 510 | 113 | 416 | not sig | 0.502 | 1.358 | 79 | 42 | 4 | 4 | -0.260 |
| p23 | 314 | 612 | 314 | 422 | not sig | 1.080 | 3.016 | 108 | 45 | 4 | 4 | -1.351 |
| p24 | 300 | 113 | 314 | 432 | not sig | -0.019 | 3.325 | 52 | 42 | 4 | 4 | 0.078 |
| p25 | 293 | 213 | 413 | 427 | not sig | 0.317 | 2.948 | 112 | 27 | 3 | 3 | -0.477 |
| p26 | 293 | 213 | 510 | 414 | not sig | -0.012 | 4.520 | 57 | 25 | 3 | 4 | 0.020 |
| p27 | 304 | 113 | 510 | 426 | not sig | 0.241 | 3.725 | 73 | 30 | 2 | 3 | -0.655 |
| p28 | 272 | 113 | 413 | 464 | sig | -0.191 | 3.134 | 45 | 21 | 3 | 4 | -0.125 |
| p29 | 314 | 213 | 314 | 425 | not sig | 0.047 | 1.694 | 122 | 23 | 2 | 2 | -0.434 |
| p30 | 296 | 113 | 510 | 405 | not sig | 0.453 | 4.241 | 74 | 47 | 3 | 4 | 0.115 |
| p31 | 317 | 413 | 213 | 401 | not sig | 0.146 | 1.365 | 34 | 23 | 3 | 3 | -0.299 |
| p32 | 208 | 213 | 510 | 301 | sig | 1.581 | 4.256 | 53 | 25 | 2 | 4 | -0.210 |
| p33 | 304 | 314 | 413 | 421 | not sig | 0.205 | 4.483 | 42 | 22 | 1 | 4 | -0.144 |
| p34 | 253 | 113 | 413 | 444 | sig | 0.423 | 4.241 | 51 | 26 | 2 | 4 | -0.326 |
| p35 | 302 | 213 | 413 | 409 | not sig | -0.445 | 3.134 | 49 | 31 | 4 | 4 | 0.115 |
| p36 | 347 | 510 | 314 | 412 | not sig | 0.594 | 4.520 | 60 | 22 | 2 | 4 | -1.018 |
| p37 | 257 | 213 | 510 | 389 | not sig | 0.200 | 2.932 | 88 | 33 | 3 | 4 | 0.237 |
| p38 | 314 | 612 | 413 | 454 | not sig | 0.536 | 4.200 | 88 | 26 | 1 | 4 | -0.757 |
| p39 | 326 | 213 | 113 | 411 | not sig | 0.271 | 2.247 | 125 | 23 | 3 | 4 | -0.790 |
| p40 | 255 | 113 | 612 | 381 | sig | 0.140 | 3.929 | 98 | 37 | 2 | 4 | -0.373 |

**CFA correction method**

Compared CFA correction methods (ICA vs rest template subtraction), using a previously pre-processed form of the dataset in this paper, with -100ms to 0ms baseline correction (in accordance with the previous preregistered pre-processing pipeline). Time windows for each cardiac event was defined as follows: R period -100ms to 100ms, T period -100 to 350ms, Diastole period 350ms to 700ms, and the total window -100 to 900ms. The ratio was calculated as the division of the sum of the root mean square across channels per time point for the CFA corrected average, by the sum of the root mean square across channels per time point for the raw data average, minus 1 (
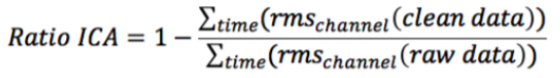
). Therefore, a higher number represents greater reduction of the CFA.

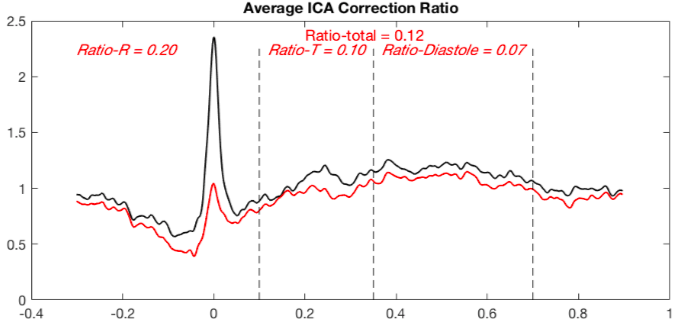

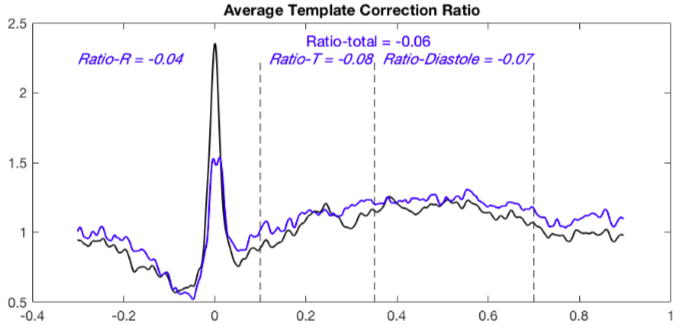

***Figure 8.*** Black line in both figures reflects averaged HEP response before CFA correction. Red line in the top figure reflects averaged HEP response after ICA CFA correction, and blue line in bottom figure reflects averaged HEP response after rest template CFA subtraction. Ratios calculated as stated above the figure.

**Control Analyses**

**HEP Control Analyses**

To ensure the R-locked main effect of delay is task-dependent and not a result of residual HEP differences, we analysed the difference between delay conditions before the first and fourth sound. We chose the fourth sound as the omission could occur from the fifth sound onward. Therefore, we computed robust averages of pre-processed HEP data relative to the R-peak for the first and fourth sound. We averaged pre-sound HEP activity belonging to the electrodes and time-window of the significant pre-omission positive cluster, for each participant. Subsequently, a two-way ANOVA analysed the interaction of cardio-audio delay (short and long delay) and sound number (first and fourth sound) and t-tests analysed the effect of cardio-audio delay separately for the first and fourth sound.

**Baseline and CFA correction controls**

To test whether the omission-locked delay effect was driven by pre-omission baseline differences (Figure 6), we performed the same comparison using the time of the significant effect (compared delay conditions 94-137ms post-omission), with two baseline correction windows (-100ms to 0ms (Pfeiffer and De Lucia., 2017; Marshall et al., 2017; 2018) and -150ms to -50ms (Sel et al., 2018; Canales-Johnson et al., 2015)). Additionally, to ensure our CFA correction method did not insert artificial effects by removing cardiac artefacts more in one condition than the other, we additionally analysed the data without CFA correction. This resulted in five control comparison combinations (-100ms to 0ms baseline correction with CFA correction, -150ms to -50ms baseline correction with CFA correction, -100ms to 0ms baseline correction with no CFA correction, -150ms to -50ms baseline correction with no CFA correction, no baseline correction with no CFA correction), in addition to the standard no baseline correction with CFA correction analyses reported throughout the paper. Equally, we completed the same five control comparisons to all ERP results which demonstrated a significant sensor level effect.

**CFA Control Analyses**

We performed control analyses on the ECG data, to determine if differences in cardiac activity contributed towards the HEP results. Therefore, we completed equivalent analyses to that which demonstrated significant ERP results on the ECG data. Subsequently, we computed single-subject robust averages of the ECG activity for each condition and analysed them using the cluster mass method, as described above. We completed ECG comparisons as to those which showed a significant ERP effect (i.e. for part 1 we compared ECG across perceived synchrony conditions time-locked to sounds 176-209ms and 240-289ms in high perceivers only. For part 2 we compared cardio-audio delay conditions 79-128ms post-R and 94-137ms post-omission, compared attention conditions 37-68ms post-R, assessed the attention and interoceptive awareness interaction 96-139ms post-omission, and assessed the pairwise effect of attention during synchronous trials in high awareness participants 105-131ms post-omission).

**Control HEP results**

We would expect a true cardio-audio expectation effect to be present not only pre-omission but also pre-sound, perhaps increasing in strength as expectation builds with the number of R-locked sounds. To test this, we analysed the main effect of delay before the fourth sound and compared this to the pre-first sound delay effect (before any sounds), using the electrodes and time window of the significant positive pre-omission cluster (R+79-128ms). Although this revealed a not significant sound number and delay interaction (F(1,33) = 0.898, *p* = .350, n^2^ = 0.007, BF_incl_ = 1.524), there was a greater delay effect before the fourth sound (Mean delay difference = 0.322), in comparison to before the first sound (Mean delay difference = 0.175).

**Baseline and CFA correction control results**

All baseline correction and CFA correction control combinations (-100 to 0, -150 to -50 and no baseline correction and with/without CFA correction) demonstrated a significant main effect of cardio-audio delay relative to the omission (largest *p* = .022), therefore CFA correction and baseline effects did not influence the omission-locked delay effect (see Supplementary Figure 9).

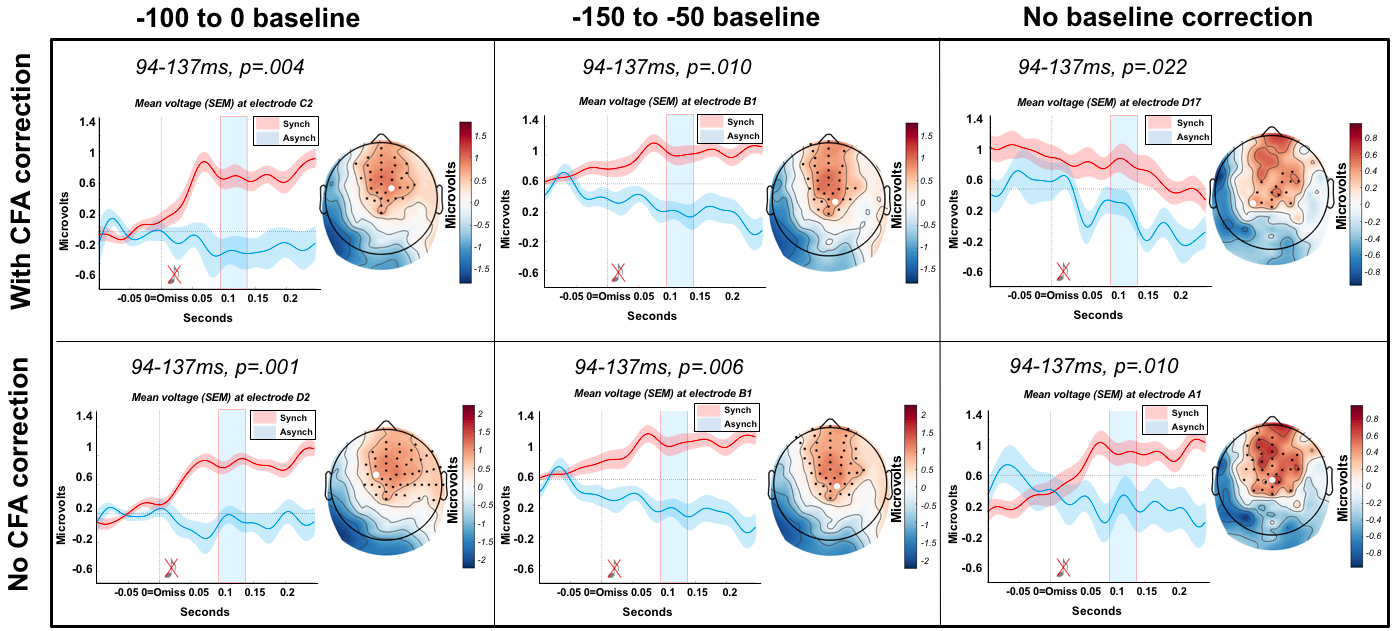

***Figure 9.*** The omission-locked delay effect with and without baseline correction (-100 to 0 and -150 to -50), as well as with and without CFA correction.

The R-locked main effect of delay was significant in all control comparisons (largest *p* = .024), although four of the control comparisons demonstrated positive and negative significant clusters rather than a sole positive cluster as demonstrated in our primary analysis results, suggesting this effect may involve broader neural regions. The R-locked main effect of attention and omission-locked attention and interoceptive awareness interaction was present in two out of the five control comparisons, with marginal significance in an additional control comparison each, suggesting that these may reflect weak effects. The 176-209ms AEP result in high perceivers was present in three and the 240-289ms AEP result was present in four out of the five control comparisons, with marginal significance in an additional control comparison for each time window (see Supplementary Table 3).

**Control ECG results**

We observed no significant differences in ECG responses between cardio-audio delay conditions 79-128ms post-R (no clusters) or 94-137ms post-omission (*p* = .128), or between attention conditions 37-68ms post-R (no clusters). Additionally, we observed no ECG differences between most rated synchronous AEPs and least rated synchronous AEPs in high perceivers, 176-209ms and 240-289ms relative to the sound (no clusters). Additionally, no significant ECG interaction of attention and awareness 96-139ms and no significant ECG simple effect of attention in high aware participants 105-131ms, post-omission and during synchronous trials only (no clusters). Therefore, it’s unlikely that ECG activity contributed towards the ERP differences observed.

**Interbeat intervals (IBI’s)**

We analysed the IBI’s throughout experimental blocks (after the removal of faulty blocks with IBI’s smaller than 400ms or larger than 1500ms). Replicating our previous study, we found that IBI’s were significantly longer when internally attending (M=838.413, SD=82.548) than when attending externally (M=827.287, SD=83.351; F(1,33) = 22.072, *p* <.001, n^2^ = 0.274, BF_incl_ = 430383.021). There was no significant IBI difference across delay conditions (F(1,33) = 0.017, *p* = .896, n^2^ = 1.026e-4, BF_incl_ = 0.157), and no significant interaction of delay and attention (F(1,33) = 0.854, *p* = .362, n^2^ = 0.003, BF_incl_ = 0.180). Thus, we can conclude that overall heart-rate differences did not influence our HEP delay effects.

Additionally, as previous studies found heart rate differences in response to omission and deviant stimuli (Banellis & Cruse, 2020; Pfeiffer & De Lucia, 2017; Raimondo et al., 2017), we investigated differences surrounding the within-task omissions. Thus, we compared the IBI around the omission (i.e. ‘omission-1’) with the following IBI (i.e. interval of the first and second R-peak after the omission ‘1-2’) and determined whether these differed between attention and delay conditions. A three-way ANOVA revealed significant effect of IBI (F(1,33) = 5.116, *p* = .030, n^2^ = .008, BF_incl_ = 0.257), a significant effect of attention (F(1,33) = 4.170, *p* = .049, n^2^ = 0.035, BF_incl_ = 4.912), a significant interaction of IBI and attention (F(1,33) = 4.555, *p* = .040, n^2^ = 0.016, BF_incl_ = 0.574), and a significant interaction of delay and attention (F(1,33) = 3.979, *p* = .054, n^2^ = 0.024, BF_incl_ = 0.812). The interaction of delay and IBI was not significant (F(1,33) = 0.125, *p* = .726, n^2^ = 1.910e-4, BF_incl_ = 0.064), and the interaction of delay, IBI and attention was not significant (F(1,33) = 0.197, *p* = .660, n^2^ = 2.942e-4, BF_incl_ = 0.045).

Posthoc tests of the IBI and attention interaction revealed a significant difference between the IBI ‘omiss-1’ during internal trials (M=841.282, SD=84.858) and IBI ‘omiss-1’ during external trials (M=831.659, SD=89.130; t(33) = 2.868, p_holm_ = .029) and a significant difference between the IBI ‘1-2’ during internal trials (M=840.152, SD=86.261) and the IBI ‘omiss-1’ during external trials (t(33) = 2.762, *p_holm_* = .033), and finally a significant difference between the IBI ‘omiss-1’ during external trials and the IBI ‘1-2’ during external trials (M=838.208, SD=90.012; t(33) = -3.030, *p_holm_* = .022). Therefore, revealing a cardiac deceleration following the omission during external trials only (as in Banellis & Cruse., 2020).

Posthoc tests of the attention and delay interaction revealed a significant difference between the IBI’s during internal perceived asynchronous delay trials and the IBI’s during external perceived asynchronous delay trials (t(33) = 2.846, *p_holm_* = .036).

**Heart rate variability**

We analysed the standard deviation of the IBI’s (SDRR) as a measure of heart rate variability. This revealed no significant HRV differences across attention (F(1,33) = 0.198, *p* = .659, n^2^ = 0.002, BF_incl_ = 0.140) or delay trials (F(1,33) = 1.133, *p* =.295, n^2^ = 0.013, BF_10_ = 0.229), as well as no significant attention and delay HRV interaction (F(1,33) = 0.536, *p* = .465, n^2^ = 0.005, BF_incl_ = 0.048), further excluding heart-related confounds.

**Multiverse controls**

**Table 3.** *Results of the baseline and CFA correction control analyses, using the significant time window of the effects with ICA CFA correction and without baseline correction (reported throughout the paper - in the black square, in bold). Green reflects significant results, although in italics if a different cluster polarity to the original result with ICA CFA correction and no baseline correction. Orange if marginally significant and red if not significant. ‘R_ME_Delay’ is the R-locked/pre-omission main effect of delay. ‘R_ME_Attention’ is the R-locked/pre-omission main effect of attention. ‘Omiss_ME_Delay’ is the omission-locked main effect of delay. ‘Omiss_Int_AttAware’ is the omission-locked interaction of attention with interoceptive awareness, with ‘Att_HighAware(SynchOnly)’ reflecting the simple/attention effect in high awareness participants and ‘Att_LowAware(SynchOnly)’ reflecting the simple/attention effect in low awareness participants only (in perceived synchronous trials only). ‘AEP_154-209_HighPerceivers’ is the first AEP effect of perceived synchrony and ‘AEP_209-289_HighPerceivers’ is the following AEP effect of perceived synchrony, both in high heartbeat perceivers only.*

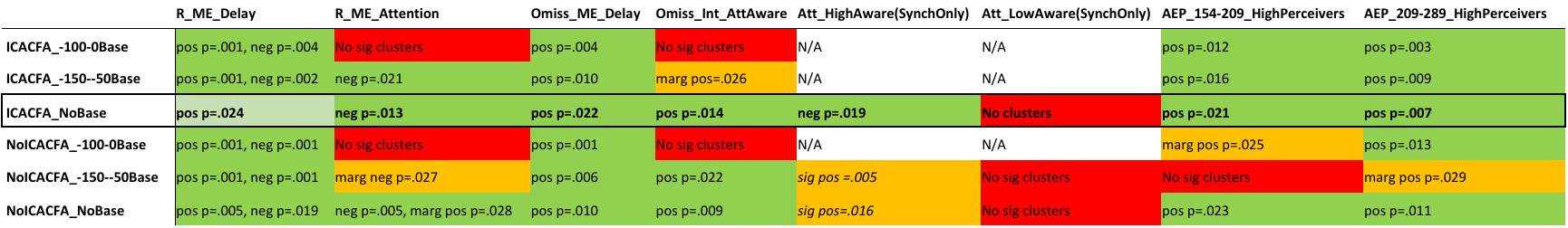
